# Previously uncharacterized interactions between the folded and intrinsically disordered domains impart asymmetric effects on UBQLN2 phase separation

**DOI:** 10.1101/2021.02.22.432116

**Authors:** Tongyin Zheng, Carlos A. Castañeda

**Affiliations:** Department of Chemistry, Syracuse University, Syracuse, NY 13244, USA; Departments of Biology and Chemistry, Syracuse University, Syracuse, NY 13244, USA; Interdisciplinary Neuroscience Program, Syracuse University, Syracuse, NY 13244, USA; BioInspired Institute, Syracuse University, Syracuse, NY 13244, USA

**Keywords:** Ubiquilin-2, liquid-liquid phase separation, Ubiquitin-like domain, Ubiquitin-associated domain, STI1 domain, intrinsically disordered regions, folded domains, self-association, NMR spectroscopy

## Abstract

Shuttle protein UBQLN2 functions in protein quality control (PQC) by binding to proteasomal receptors and ubiquitinated substrates via its N-terminal ubiquitin-like (UBL) and C-terminal ubiquitin-associated (UBA) domains, respectively. Between these two folded domains are intrinsically disordered STI1-I and STI1-II regions, connected by disordered linkers. The STI1 regions bind other components, such as HSP70, that are important to the PQC functions of UBQLN2. We recently determined that the STI1-II region enables UBQLN2 to undergo liquid-liquid phase separation (LLPS) to form liquid droplets *in vitro* and biomolecular condensates in cells. However, how the interplay between the folded (UBL/UBA) domains and the intrinsically-disordered regions mediates phase separation is largely unknown. Using engineered domain deletion constructs, we found that removing the UBA domain inhibits UBQLN2 LLPS while removing the UBL domain enhances LLPS, suggesting that UBA and UBL domains contribute asymmetrically in modulating UBQLN2 LLPS. To explain these differential effects, we interrogated the interactions that involve the UBA and UBL domains across the entire UBQLN2 molecule using NMR spectroscopy. To our surprise, aside from well-studied canonical UBL:UBA interactions, there also exist moderate and weak interactions between the UBL and STI1-I/STI1-II domains, and between the UBA domain and the linker connecting the two STI1 regions, respectively. Our findings are essential for the understanding of both the molecular driving forces of UBQLN2 LLPS and the effects of ligand binding to UBL, UBA, or STI1 domains on the phase behavior and physiological functions of UBQLN2.

**Impact of Work Statement:** Zheng and Castañeda show that interplay between the folded domains and intrinsically disordered regions regulates liquid-liquid phase separation behavior of UBQLN2, a protein quality control (PQC) shuttle protein. Despite their similar size, the folded UBL and UBA domains inhibit and promote phase separation, respectively, due to their previously uncharacterized, asymmetric interactions with the middle intrinsically-disordered region. These results strongly suggest that PQC components, including proteasomal receptors, are likely to modulate UBQLN2 phase separation behavior in cells.

## Introduction

Protein degradation by the ubiquitin-proteasome system (UPS) involves recognition of ubiquitinated substrates by receptors located on the proteasomal regulatory particle or extraproteasomal receptors called ubiquitin (Ub)-binding shuttle proteins (Martinez-Fonts et al., 2020; Zientara-Rytter and Subramani, 2019). UPS shuttle proteins are defined by a UBL-UBA domain architecture, whereby the UBL (Ubiquitin-Like) domain interacts with proteasomal receptors and the UBA (Ubiquitin-associated) domain interacts with ubiquitin chains on ubiquitinated substrates marked for degradation (Chen et al., 2019; Ko et al., 2004). One such shuttle protein is ubiquilin-2 (UBQLN2), which undergoes liquid-liquid phase separation and is recruited to cytoplasmic stress granules, biomolecular condensates with liquid-like properties (Alexander et al., 2018; Dao et al., 2018). As Ub binds to UBQLN2 UBA to drive disassembly of UBQLN2 condensates, we hypothesize that LLPS contributes to UBQLN2’s functionality in UPS and protein quality control (PQC) (Dao and Castañeda, 2020; Zheng et al., 2020). Recently, other UBL-UBA shuttle proteins including hHR23B and p62/SQSTM1 have been reported to undergo LLPS in the presence of polyUb chains (Sun et al., 2018; Yasuda et al., 2020; Zaffagnini et al., 2018). Aside from *in vitro* evidence that these shuttle proteins phase separate, hHR23B and SQSTM1 colocalize with nuclear proteasomal foci and autophagosome precursors, respectively. Therefore, it is critical to delineate the contributions of the UBL and UBA domains to LLPS to understand the role of phase separation to the functionality of shuttle proteins.

UBQLN2 is a 624-amino acid multidomain protein composed of an intrinsically-disordered region (IDR) capped by N-terminal UBL and C-terminal UBA domains (Figure 1A). Since UBL and UBA domains are common features of many PQC proteins, their structures have been studied extensively (Chen et al., 2001, 2019; Díaz-Martínez et al., 2006; Heir et al., 2006; Ohno et al., 2005; Walters et al., 2002; Zhang et al., 2008). The middle IDR of UBQLN2, consisting of residues 109-576, includes the STI1-I, STI1-II and PXX regions connected by disordered linkers. The STI1 regions are presumed to mediate interactions with chaperone proteins such as HSP70, autophagosomal components such as LC3, and even client proteins (Hjerpe et al., 2016; Kurlawala et al., 2017; Lee et al., 2013; Zheng et al., 2020). However, specific interactions involving the IDR are largely unknown.

**Figure 1.**
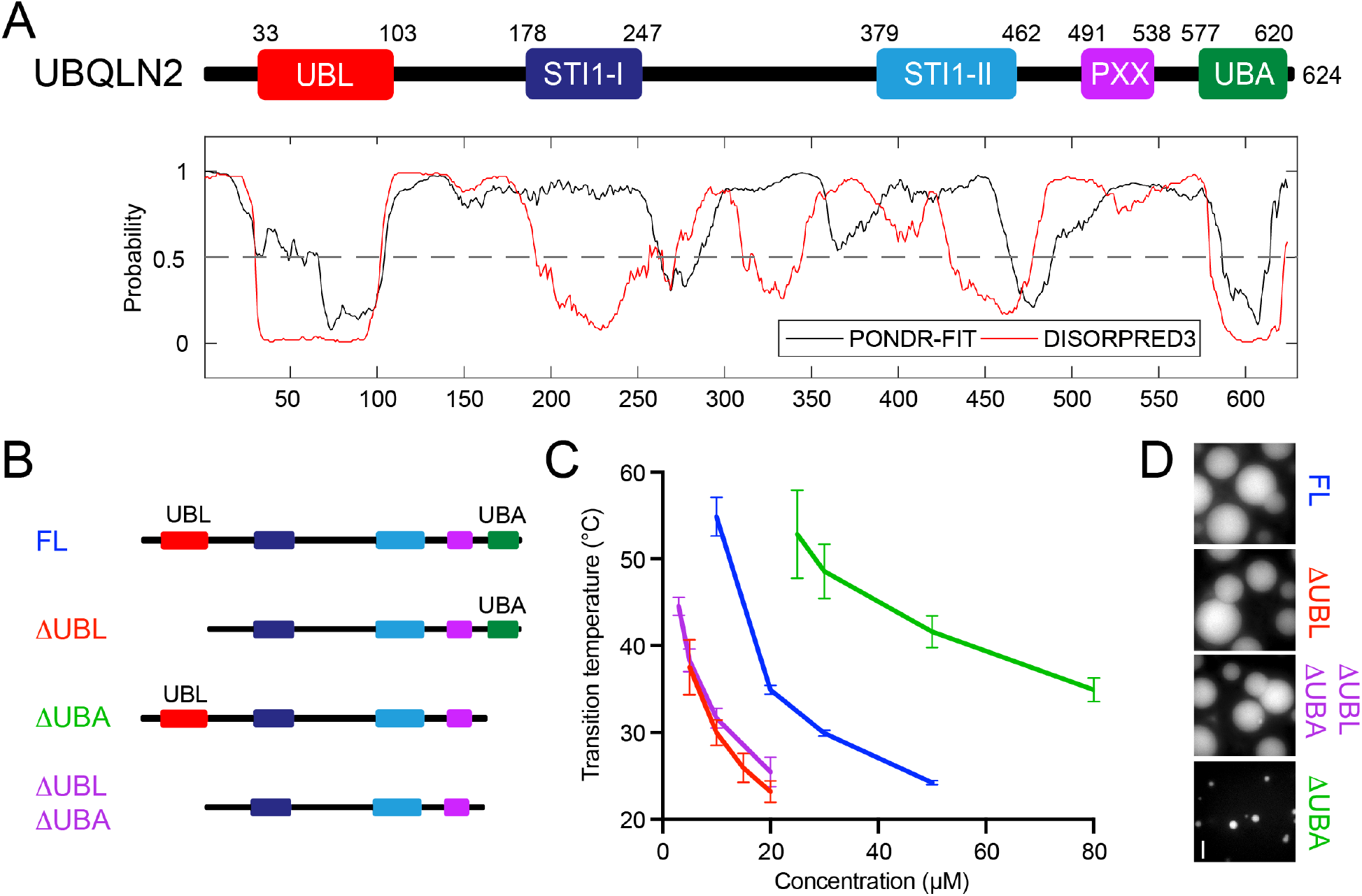
UBL and UBA domains contribute asymmetrically to UBQLN2 LLPS. (A) Domain architecture of UBQLN2. Disordered propensity predicted using the program PONDR-FIT (Xue et al., 2010), and DISORPRED3 (Jones and Cozzetto, 2015). (B) Library of UBL and UBA domain deletion constructs used here and color-coded accordingly. (C) Temperature-concentration phase diagram constructed from spectrophotometric assays using denoted protein concentrations in pH 6.8 buffer containing 200 mM NaCl. Error bars are standard deviation from at least four measurements. (D) Fluorescence microscopy shows solutions of UBQLN2 domain deletion constructs under identical conditions (100 μM protein, spiked with Dylight-488 labelled FL-UBQLN2 protein at 1:100 molar ratio, pH 6.8 20 mM NaPhosphate, 0.5 mM EDTA, 0.02% NaN_3_, 200 mM NaCl, at 30 °C). Scale bar denotes 5 μm.

Our lab previously found the STI1-II region to be essential for the oligomerization and LLPS of UBQLN2. Constructs that lack the STI1-II regions do not oligomerize or undergo LLPS under physiological conditions (Dao et al., 2018). On a molecular level, we used a C-terminal construct of residues 450-624 to show that multiple transient interactions exist among the STI1-II, PXX and UBA domains (Dao et al., 2018, 2019) to collectively drive UBQLN2 self-association and phase separation. Ubiquitin-binding to the UBA significantly reduces the driving forces for UBQLN2 phase separation by decreasing the number of multivalent “stickers” needed for LLPS, thus driving the disassembly of UBQLN2 condensates *in vitro*. Importantly, it is well-established that UBL and UBA domains interact with each other (Lowe et al., 2006; Nguyen et al., 2017). How interplay among these folded domains and the middle region of UBQLN2 mediates phase separation remains poorly understood.

In this study, we systematically mapped out the interactions that occur between the folded UBL, UBA domains and the middle parts of the UBQLN2 protein using NMR spectroscopy and a library of domain deletion constructs, and determined how these interactions modulate UBQLN2 phase separation. Unexpectedly, our results indicated that the UBL and UBA domains interact with separate parts of 109-576. We uncovered moderate and specific interactions between UBL and each STI region, and weak interactions between the UBA domain and linker residues 248-378. Importantly, we find that these interdomain interactions have opposing effects on the LLPS behavior of UBQLN2. Deletion of the UBL domain enhances the driving forces for LLPS, whereas deletion of the UBA domain inhibits LLPS. Our results suggest that the IDR-driven LLPS behavior of UBQLN2 can be tuned by ligand interactions involving either the UBL or UBA domains. Given that UBQLN2’s function as a shuttle protein heavily relies on interactions involving the UBL and UBA domains, our data support the hypothesis that LLPS contributes to the physiological functions of UBQLN2.

## Results

### UBQLN2 LLPS does not require UBL or UBA domains for phase separation

To determine the contribution of the UBL and UBA domains to UBQLN2 LLPS, we generated several UBQLN2 constructs with the UBL and/or UBA domains removed: ΔUBL (residues 109-624), ΔUBA (1-576), and ΔUBLΔUBA (109-576) (Figure 1B). We expressed and purified these constructs in *E. coli* (see Methods, Figure S1)). Importantly, all constructs phase separated with increasing salt and temperature, consistent with full-length (FL) UBQLN2 phase behavior. Thus, we were able to purify the proteins without any extra mutations or affinity tags (Dao et al., 2018). We obtained the low-concentration arm of the temperature-concentration phase diagram for each UBQLN2 construct using spectrophotometric turbidity assays performed in 20 mM NaPhosphate buffer with 200 mM NaCl, pH 6.8 (Figure S2). The low-concentration phase boundary denotes the saturation concentration (c_sat_) above which the protein phase separates (Figure 1C). Using fluorescence microscopy, we confirmed that each construct formed protein-rich, liquid-like condensates (Figure 1D, Movie S1-S4). Importantly, our data showed that UBQLN2 does not require the folded UBL or UBA domains to phase separate, consistent with prior data from our group and others (Alexander et al., 2018; Dao et al., 2018).

### UBL and UBA domains contribute asymmetrically to UBQLN2 LLPS

Removal of the UBL domain (ΔUBL) and UBA domain (ΔUBA) from FL UBQLN2 had opposite effects on the phase behavior of UBQLN2 (Figure 1C). Over the 16-60 °.C temperature range, phase separation of ΔUBA required much higher protein concentrations (high c_sat_ values) compared to FL UBQLN2, whereas ΔUBL phase-separated at much lower protein concentrations (low c_sat_ values). The effect of removing both UBL and UBA domains was not additive, as the phase diagram of 109-576 was nearly identical to that of ΔUBL. From these observations, we hypothesized that the asymmetrical contributions of UBL and UBA domains on UBQLN2 LLPS reflect differences in how these folded domains interact with the rest of the UBQLN2 protein. To parse these interactions, we constructed a domain deletion library of UBQLN2 constructs, and employed NMR spectroscopy to map the interactions involving the folded UBL and UBA domains across the entire UBQLN2 protein.

### UBQLN2 UBL and UBA domains interact with one another

It is well-established that UBL and UBA domains of Ub-binding shuttle proteins interact with each other on an intra- and intermolecular basis (Lee et al., 2013; Lowe et al., 2006; Nguyen et al., 2017). To validate this within our UBQLN2 constructs, we monitored the interactions between the isolated UBA and UBL domains of UBQLN2 using NMR spectroscopy (Figure 2), similar to the experiments employed in (Nguyen et al., 2017). We used UBQLN2 1-107 and UBQLN2 571-624 as our UBL and UBA constructs, respectively, with a C-terminal His-tag (see Methods and Table S1). We obtained chemical shift assignments for the UBL construct, and found that the N-terminal 30 residues are disordered, consistent with secondary structure predictions (Figure 1A). As a control, we compared the spectra of the UBL or UBA domains at 50 and 400 μM and observed minimal or no concentration-dependent chemical shift perturbations (CSPs) (Figure S3), which suggest that these constructs do not self-associate over this concentration range.

**Figure 2.**
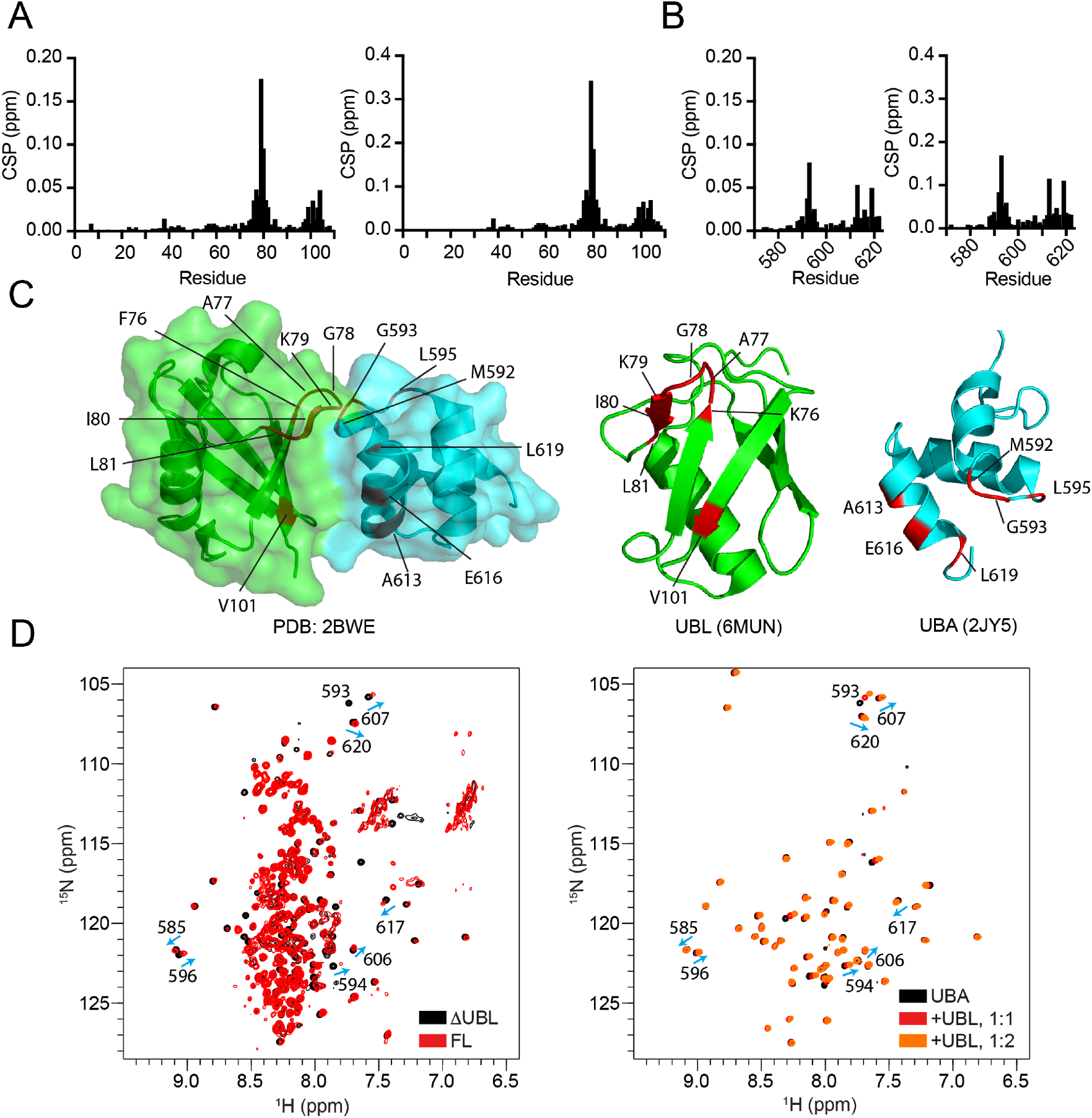
UBQLN2 UBA and UBL domains interact with each other. >(A, B) Chemical shift perturbations (CSPs) of ^1^H-^15^N amide resonances of (A) the isolated UBL domain (left: 50 μM, right: 200 μM) in the presence of equimolar amounts of UBA domain and (B) the isolated UBA domain (left: 50 μM, right: 200 μM) in the presence of equimolar amounts of UBL domain. (C) Mapping of UBQLN2 UBL and UBA CSPs on the UBL:UBA binding surface of yeast ubiquilin homolog DSK2 (PDB ID: 2BWE). UBL residues with CSP > 0.03 ppm and UBA residues with CSP > 0.02 ppm from panel A are highlighted in red; structures of isolated UBQLN UBL and UBA are shown on the right (PDB ID: 6MUN, PDB ID: 2JY5). (D) Overlays of (left) FL UBQLN2 and ti. UBL spectra and of (right) isolated UBA and UBA:UBL spectra. All proteins at 50 μM concentration except for UBA:UBL at 1:2 molar ratio where UBL is 100 μM. UBA peaks in FL UBQLN2 are similarly affected by inter- and intra-molecular UBL binding as peaks in isolated UBA spectrum in the presence of isolated UBL domain.

We mapped the UBL:UBA binding interface by identifying those residues that exhibited significant CSPs upon the titration of one folded ^15^N-labeled domain with the other (unlabeled) domain. Consistent with other studies (Lowe et al., 2006; Nguyen et al., 2017), we observed concentration-independent backbone amide CSPs for UBL residues 75-82 and 98-103 and UBA residues 591-594 and 612-620 (Figures 2A,B). Together, these residues form a hydrophobic interface across the β sheets on the UBL and the loop 1 and a3 helix on the UBA domain (Figures 2C). These key amino acids at the UBL:UBA interface are also essential for UBL:proteasome subunit (Rpn10, Rpn13) (Chen et al., 2016, 2019), and UBA:ubiquitin interactions (Ohno et al., 2005; Zhang et al., 2008). By following chemical shift trajectories for amide resonances on both domains, we determined binding dissociation constants (K_d_) of 178 +/− 41 μM and 210 +/− 15 μM when monitoring UBL and UBA resonances, respectively (Figure S4). Our experiments are consistent with previous studies showing that UBA and UBL domains of UBQLN2 interact with weak affinity (K_d_ ~ 200 μM) (Nguyen et al., 2017).

To test whether UBL-UBA interactions are present in the intact FL UBQLN2 protein, we collected TROSY-HSQC spectra of FL UBQLN2 and ΔUBL. While we didn’t see UBL resonances in FL UBQLN2 spectra, UBA resonances were clearly visible (Figure 2D and (Dao et al., 2018)). The UBA peaks in ΔUBL spectra superpositioned well with those in isolated UBA spectra, whereas UBA peaks in FL UBQLN2 approached equimolar UBA:UBL peak positions, suggesting that UBA is interacting with the UBL within or between UBQLN2 molecules in a similar manner as in the isolated UBA:UBL experiments (Figure 2D).

### UBL interacts with the middle STI1 domains of UBQLN2

Given the asymmetric contributions of the UBL and UBA domains to the phase behavior of UBQLN2, we suspected that the UBL and UBA domains differentially interacted with the intrinsically-disordered middle region of UBQLN2 (residues 109-576). To probe these interactions, we first compared ^1^H-^15^N NMR spectra of 200 μM ^15^N UBL domain in the absence and presence of equimolar amounts of unlabeled UBQLN2 109-576. All NMR experiments were conducted under conditions where the proteins did not phase separate (no added NaCl). When 109-576 was added to UBL, we noticed that nearly all the UBL peaks broadened beyond detection, with the exception of the flexible N-terminal residues 1-30 (Figure 3A). The strong attenuation of UBL peaks indicated that UBL binds to the middle region of UBQLN2, which excludes the folded UBA domain, and also provides an explanation for why UBL peaks are not visible in FL UBQLN2 spectra.

**Figure 3.**
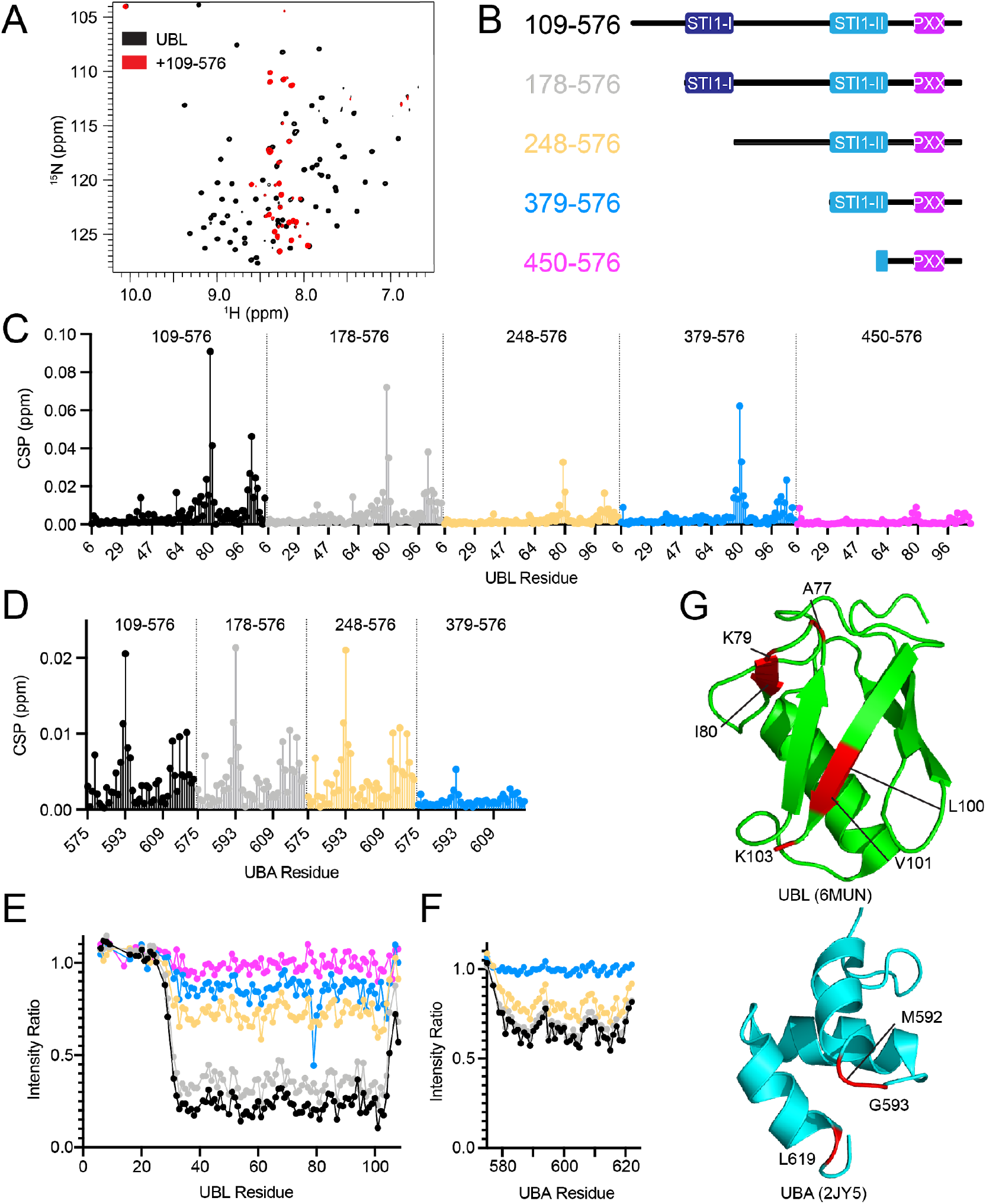
UBL and UBA domains interact with distinct regions in the intrinsically disordered segment of UBQLN2. (A) UBL NMR signals attenuated upon addition of UBQLN2 109-576. (B) Library of domain deletion constructs designed to probe UBL/UBA:109-576 interactions. (C) CSPs of UBQLN2 UBL upon mixing 50 μM UBL with equimolar amount of designated UBQLN2 domain deletion construct, or (D) CSPs of UBQLN2 UBA domain upon mixing 200 μM UBA with equimolar amount of designated UBQLN2 domain deletion construct. (E, F) Amide peak intensity ratios of resonances in either (E) UBL or (F) UBA in the presence of designated UBQLN2 constructs. (G) CSPs arising from UBL/UBA interactions with 109-576. CSPs (red) are mapped onto UBL and UBA structures: CSPs > 0.02 ppm and CSPs > 0.01 ppm for UBL and UBA, respectively.

We speculated that UBL peak broadening stemmed from slow tumbling of the UBL:109-576 complex as 109-576 contains the oligomerization-driving STI1-II region (Dao et al., 2018). We reasoned that lowering 109-576 concentration would reduce oligomerization and enable the tracking of UBL resonances upon binding. When we repeated our NMR experiments using ^15^N 50 μM UBL and added equimolar amounts of 109-576, UBL peaks remained visible, even though we observed a significant reduction (up to 80%) in amide peak intensity (Figure 3). Our CSP data revealed that the UBL residues that interact with 109-576 are identical to those that interact with the UBA domain, specifically UBL residues 77-80 and 99-101. Notably, the UBL:109-576 interactions are specific to the UBL as Ub (identical fold but different sequence composition) did not interact with 109-576 (Figure S5A).

The UBL:109-576 CSPs were reduced in half compared to the UBL:UBA CSPs under identical conditions. To determine the UBL binding affinity for 109-576, we attempted a NMR titration experiment where we incrementally added 109-576 to 50 μM UBL. However, UBL signals attenuated beyond detection when > 70 μM 109-576 was added (Figure S5B), likely owing to the propensity for 109-576 to oligomerize and form a UBL:109-576 complex that tumbles too slowly for NMR detection. Notably, the peak trajectories for UBL resonances are not in the same direction when comparing the UBL:UBA and UBL:109-576 titration experiments, indicative of differences in binding modes. Based on the totality of our data, we surmise that the UBL:109-576 affinity is of the same order of magnitude but weaker than UBL:UBA interactions.

To probe which part of the 109-576 sequence interacts with the UBL domain, we built a library of deletion constructs from the N-terminus of 109-576 (Figure 3B). These four constructs are 178-576 (deletion of linker between UBL and STI1-I region), 248-576 (subsequent deletion of STI1-I), 379-576 (subsequent deletion of linker between STI1 regions), 450-576 (subsequent deletion of most of STI1-II). Using 50 μM UBL as our reference spectrum, we mixed equimolar amounts of these constructs one at a time, and measured CSPs and peak intensities (Figures 3C and 3E). First, CSPs and peak intensities were largely comparable when UBL was mixed with either 109-576 or 178-576, suggesting that linker residues 109-177 do not interact with UBL. Strikingly, we saw a significant decrease in UBL CSPs when the entire STI1-I region was removed (compare 178-576 to 248-576), and similarly again when STI1-II residues 379-450 were removed (compare 379-576 to 450-576). Furthermore, UBL peak intensities were restored to 70% and 100% of levels for the isolated UBL when the STI1-I and most of STI1-II regions were removed, respectively. Together, these data suggested that UBL interacts with both STI1 regions. Interestingly, UBL CSPs in the presence of 379-576 were higher than with 248-576. We speculate that linker residues 248-379 may inhibit UBL access to the STI1-II region, as the 379-576 construct interacts with UBL with similar CSPs as the 178-576 construct (Figure 3C). Our deletion construct data are consistent with a model whereby UBL interacts with the STI1-I and STI1-II domains of UBQLN2, but not the interdomain linkers.

To monitor the UBL:109-576 interaction from the 109-576 side, we prepared a ^15^N-labeled sample of 109-576. Initial spectral characterization revealed that only ~160 amide resonances (out of an expected 400) were present. Overlay with our previously-characterized UBQLN2 450-624 indicated that the resonances for the disordered PXX and linker regions (residues 510-550) superimposed well, but resonances for residues 450-480 in the C-terminal region of the STI1-II were not observable. Addition of the UBL resulted in minimal or no changes to the spectra of 109-576, both in terms of peak positions and intensities (Figure S6). In combination with our data above, we concluded that the STI1-I and STI1-II resonances in the 109-576 construct are not observable via NMR, either due to high oligomerization propensity or other dynamics.

### UBA interacts very weakly with the middle IDR region of UBQLN2

To determine whether the UBA domain interacts with 109-576, we performed NMR titration experiments where we used 200 μM ^15^N-labeled UBA, and added an equimolar amount of 109-576. Unlike with UBL, UBA peaks were well-resolved in the presence of 109-576, and UBA peak intensities were only slightly attenuated (60% of their levels) (Figure 3F). In sharp contrast to what we observed with UBL, the UBA CSPs in the presence of 109-576 were significantly weaker, as the maximum UBA CSP was only 0.02 ppm whereas the maximum UBL CSP was 0.10 ppm even though we used four times less protein for the UBL experiments (Figure 3D). The UBA CSPs map to the same hydrophobic binding surface involved in interactions with either the UBL domain or with ubiquitin (Tse et al., 2011; Zhang et al., 2008). Largest CSPs (≤ 0.02 ppm) are observed for residues 592, 593, 616, and 619. These experiments indicate that the UBA interacts much more weakly with the 109-576 region of UBQLN2 than the UBL domain.

To map out the segment within 109-576 that interacts with the UBA domain, we mixed equimolar amounts (200 μM) of the UBA domain with our library of 109-576-derived constructs. We observed identical overall CSP patterns and peak intensities for UBA resonances in the presence of 109-576, 178-576, or 248-576 constructs (Figure 3D). However, nearly no UBA CSPs (all CSPs < 0.005 ppm) were observed with the equimolar UBA:379-576 mixture and UBA peak intensities were largely identical to that of the isolated UBA domain. These results suggested weak interactions between the UBA and linker residues 248-378. Additionally, the data suggested that the UBA domain minimally interacts with residues 379-576, consistent with our prior paramagnetic relaxation enhancement (PRE) data where we saw transient interactions among the UBA and C-terminal end of STI1-II region (Dao et al., 2019).

### Molecular mechanisms underlying UBL/UBA effects on UBQLN2 LLPS

Given the differences in interaction strength and binding site preferences between the UBL/UBA domains with 109-576, we rationalized that titration of the UBL and UBA domains would exert different effects on UBQLN2 109-576 LLPS. To test this, we performed our turbidity assays with 10 μM 109-576 and different amounts of isolated UBL or UBA (Figure 4A). Addition of UBL steadily increased the phase transition midpoint temperature, suggesting a lower propensity for the UBL:109-576 mixture to phase separate. This result is consistent with the phase diagram of ΔUBA (1-576) shifting to higher temperatures and concentrations than FL UBQLN2. In contrast, titration of isolated UBA to 109-576 only marginally affected UBQLN2 phase separation, suggesting that UBA:109-576 interactions are weaker than UBL:109-576 interactions in agreement with our NMR data above (Figure 4A, 4B). Notably, the effect of UBL/UBA titrations is small, likely due to the low concentration of 109-576 used (10 μM) and because we titrated UBL/UBA in *trans*.

**Figure 4.**
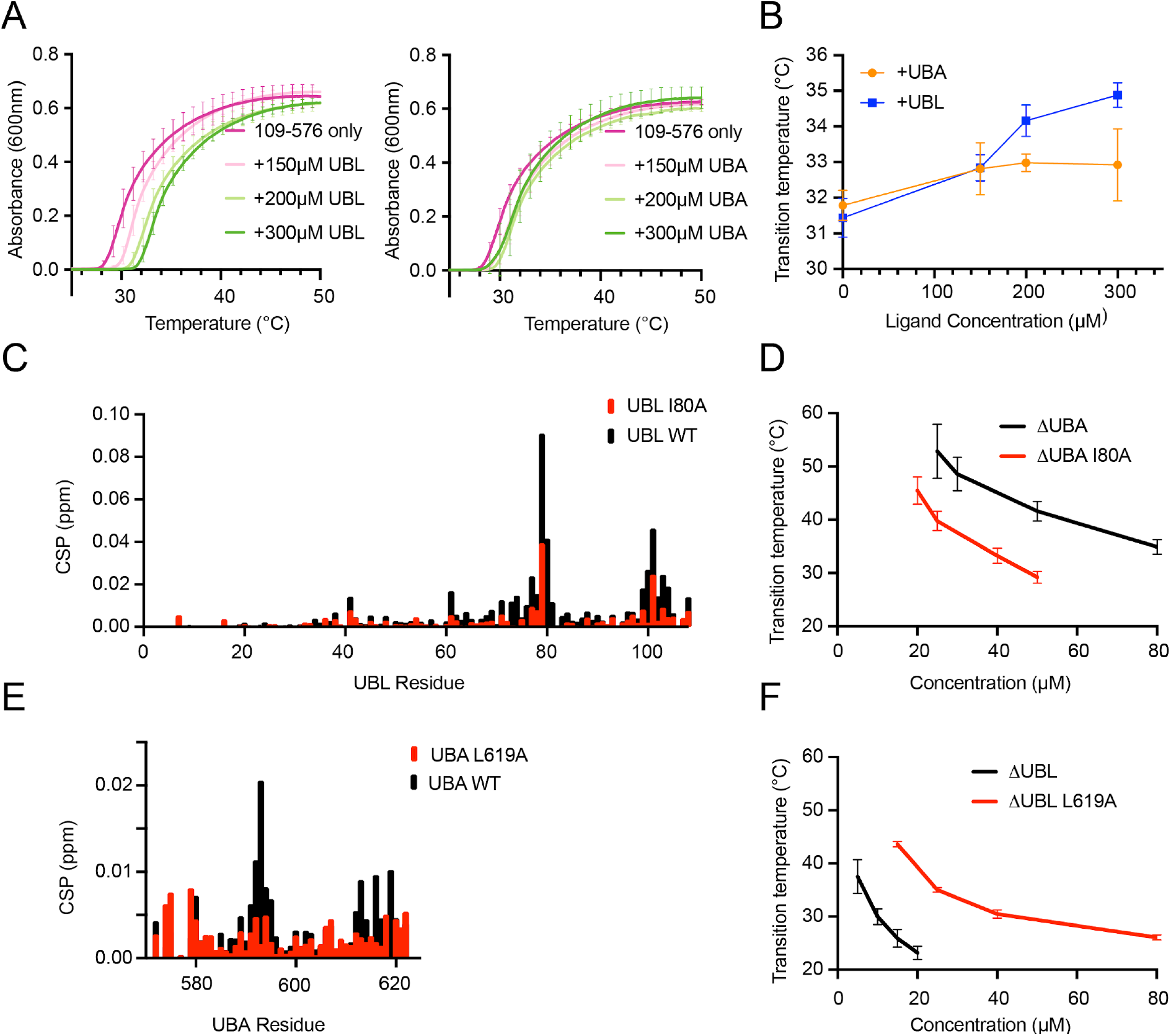
UBL:109-576 and UBA:109-576 interactions contribute asymmetrically to UBQLN2 LLPS. (A) Turbidity curves as UBL or UBA were titrated into solutions of 10 μM 109-576. (B) Plot of the changes in UBQLN2 109-576 transition temperature from panel A as UBL or UBA were titrated. (C) CSPs of UBQLN2 UBL upon mixing 50 μM UBL with equimolar amount of 109-576 for WT UBL (black) and UBL I80A (red). (D) Temperature-concentration phase diagram indicating UBL I80A mutation promotes UBQLN2 ΔUBA LLPS. (E) CSPs of UBQLN2 UBA upon mixing 200 μM UBA with an equimolar amount of 109-576 for WT UBA (black) and UBA L619A (red). (F) Temperature-concentration phase diagram indicating UBA L619A mutation inhibits UBQLN2 ΔUBL LLPS.

Our ΔUBL LLPS data suggest that the UBL “inhibits” UBQLN2 LLPS, as removal of the UBL domain lowers the protein c_sat_ threshold for phase separation. As STI1-II is critical for UBQLN2 self-association and phase separation (Dao et al., 2018) and our NMR data indicate that UBL interacts with both STI1 regions in UBQLN2, we hypothesized that UBL:STI1 interactions compete with STI1-driven self-association to abrogate phase separation. To test this hypothesis, we introduced a mutation (I80A) in the UBL domain that weakens the UBL:STI1 interactions but doesn’t impact the overall UBL structure (Figure 4C, Figure S7A). NMR experiments of I80A UBL with 109-576 confirmed that CSPs and peak intensity attenuations are smaller than those for WT UBL, indicative of weaker binding between I80A UBL and 109-576. To determine the effects of weakened UBL:STI1 interactions on LLPS, we generated I80A ΔUBA. We chose the ΔUBA background to eliminate possible UBL:UBA contributions. We predicted that STI1-driven interactions outcompete weakened UBL:STI1 interactions to promote I80A ΔUBA LLPS over a greater temperature and concentration range relative to ΔUBA. We observed this exact trend, suggesting that the UBL negatively regulates UBQLN2 LLPS (Figure 4D).

Finally, we investigated the contributions of UBA:109-576 interactions on UBQLN2 LLPS. We introduced the L619A mutation in the UBA domain and determined that L619A reduces the interactions between UBA and 109-576 using NMR spectroscopy (Figure 4E). We expressed and purified L619A ΔUBL and obtained a phase diagram to compare against ΔUBL. We found that L619A ΔUBL phase separated at higher c_sat_ values than ΔUBL (Figure 4F). These data suggest that the wild-type UBA domain favorably contributes to UBQLN2 LLPS. Given the weak UBA:109-576 interactions, we were surprised at the extent to which L619A significantly decreased LLPS propensity of ΔUBL. However, residue 619 is considered a “sticker” for LLPS, therefore the L619A mutation may also act to reduce phase separation by reducing the multivalency (number of “stickers”) necessary for LLPS (Yang et al., 2019).

## Discussion

Here we showed that the two folded domains of UBQLN2 asymmetrically modulate its IDR-driven phase separation behavior. In elucidating the molecular basis for these observations, we postulate that differences in interaction strengths and previously uncharacterized binding site preferences between the folded domains and the middle region of UBQLN2 (109-576) establish a hierarchy of interactions, some of which compete with known STI1 interactions that promote phase separation of UBQLN2. We have experimentally mapped and rank-ordered the interactions involving the folded UBL and UBA domains across UBQLN2, specifically UBL:UBA, UBL:109-576 and UBA:109-576 interactions (Figure 5A). Our NMR data suggest that UBA:109-576 interactions are the weakest of the three, while UBL:109-576 and UBL:UBA are of the same order of magnitude. UBL:UBA interactions are the strongest as they exhibit the highest CSPs among the three sets presented here. While our experiments assessed these interactions in *trans* (intermolecular interactions between designed UBQLN2 constructs), we expect these interactions to be strengthened in *cis* due to the domains being tethered together, effectively increasing their local concentrations (Kjaergaard et al., 2021).

**Figure 5.**
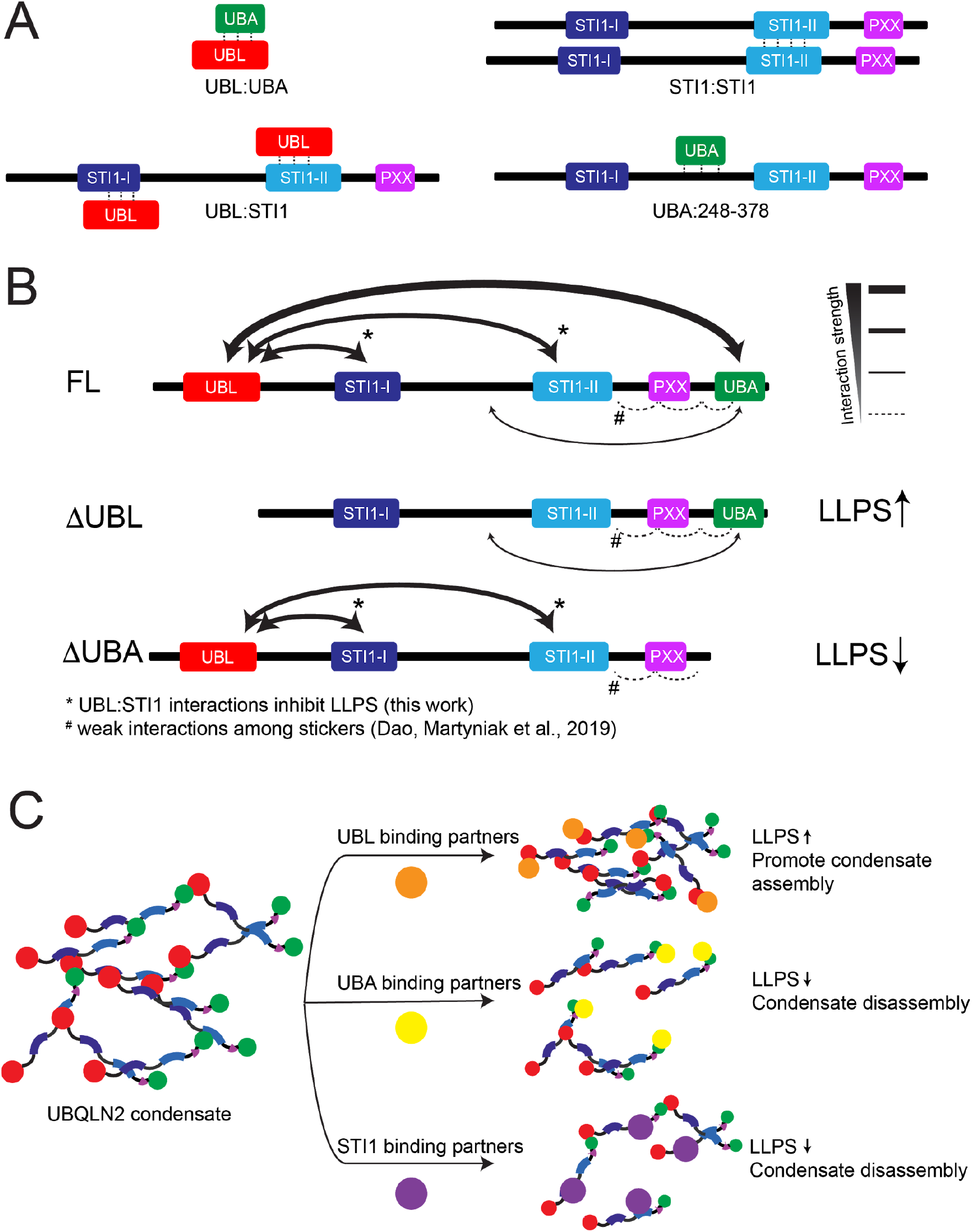
Implications of UBL/UBA interactions on UBQLN2 LLPS. (A) Summary of domain-specific interactions across UBQLN2. (B) Estimated domain-domain interactions and their interaction strengths (arrow thickness) in different UBQLN2 constructs. Interaction strengths estimated from NMR measurements (this work unless noted). (C) UBQLN2 binding partners likely modulate the driving forces of UBQLN2 phase separation depending on where they bind to UBQLN2. As drawn, the model assumes binding partners are monovalent or exhibit low valency. Throughout the figure, UBQLN2 domains are colored according to Figure 1A.

### UBQLN2 LLPS is modulated by its interdomain interaction network

The intermediate phase behavior of FL compared to ΔUBL and ΔUBA can be explained by the complex interplay of UBL:UBA, UBL:109-576 and UBA:109-576 interactions (Figure 5A). When only the UBA domain is removed from FL (ΔUBA construct, which has the lowest driving forces for phase separation among the three), two LLPS-driving “sticker” regions (UBA residues 592-594, 616-620) are eliminated, thus reducing the multivalency required for phase separation (Dao et al., 2018). Additionally, removing UBA eliminates competing UBL:UBA interactions and enables UBL to interact with the STI1 regions, thus further interfering with STI1-mediated oligomerization and phase separation. These two effects combine to reduce phase separation of ΔUBA compared to FL (Figure 5B). Conversely, phase separation of ΔUBL is promoted compared to FL. This behavior stems from at least two contributions: (1) elimination of UBL:STI1 interactions that would compete with STI1-mediated LLPS, and (2) release of UBA from UBL:UBA interactions enables UBA’s participation in LLPS-mediating interactions. Unlike UBL, UBA does not compete with STI1-mediated LLPS as there are minimal UBA:STI1 interactions. Together, these observations suggest that, in FL UBQLN2, the UBL:UBA interactions simultaneously enhance and inhibit UBQLN2 LLPS by outcompeting UBL:STI1 interactions to free the STI1 regions to drive LLPS and by reducing the availability of UBA “stickers” to participate in LLPS, respectively (Figure 5B).

Strikingly, the low concentration arms of the ΔUBL and 109-576 (ΔUBLΔUBA) phase diagrams are almost superimposable (Figure 1C). As removal of the UBA domain takes away some of the multivalent stickers required for UBQLN2 LLPS, we would predict decreased LLPS of 109-576 in comparison to ΔUBL. However, 109-576 is predicted to be largely disordered (Figure 1A), whereas ΔUBL still contains the folded UBA domain. Deletion of both UBL and UBA domains could enhance the ability of the middle intrinsically-disordered region (IDR) to interact with itself, thus promoting LLPS (Zhou et al., 2018). These two countering effects could cancel each other to yield similar phase diagrams for 109-576 and ΔUBL.

### Molecular basis of UBL:STI1 interactions

For the first time, we identified that the UBL domain interacts with the middle STI1 regions of UBQLN2. The UBL interaction patch in the UBL:STI1 interface includes the same hydrophobic residues that are in the UBL:UBA interface, specifically V73, L74, F76, A77, I80, L81, L100, V101, and I102, as well as basic residues K79 and K103. While no structural information yet exists for the STI1 regions, sequence analysis reveals that both STI1s are enriched with hydrophobic (e.g. L, M, P) and polar (N, Q) residues (Figure S8). Notably, the UBL domain is basic (pI ~ 9.4), whereas the rest of the protein is acidic (pI of 4.5 for 109-576, pI of 4.6 for UBA, see Table S5). If favorable electrostatics contributed to UBL:STI1 and UBL:UBA interactions, we would have expected UBL CSPs to be similar across all 109-576 constructs as all have identical pI values, but this was not the case (Figure 3C). Electrostatics contributions are expected to be small given the low number of ionizable residues across UBQLN2. Therefore, we speculate that the UBL:STI1 interactions largely involve hydrophobic and non-ionizable residues. However, other segments in UBQLN2 109-576 are also enriched in hydrophobic and polar residues (e.g. linker between the two STI1 regions, Figure S8), but lack significant interaction with UBL. This suggests that specific UBL-binding sites in the STI1 regions exist.

### Implications for how PQC components modulate UBQLN2 phase transitions

Our results imply that UBL- and UBA-mediated interactions modulate UBQLN2 phase transitions in cells (Figure 5C). We envision that ΔUBL mimics the UBL-bound state, e.g. when UBQLN2 UBL is bound to the proteasome via receptors such as Rpn10 or Rpn13. In such a scenario, our model predicts that UBL:receptor interactions (*K*_d_ < 10 μM) (Chen et al., 2019) outcompete both UBL:STI1 and UBL:UBA interactions, resulting in more available STI1 and UBA regions, respectively, to facilitate LLPS. On the other hand, the ΔUBA construct mimics when UBA interacts with Ub on ubiquitinated substrates, making UBA unavailable to contribute to UBQLN2 LLPS. Strong UBA:Ub interactions (*K*_d_ ~ 1-5 μM) outcompete UBA:UBL ones, thus permitting unbound UBL to engage with the STI1 regions and compete with STI1-mediated interactions for LLPS. Consequently, UBQLN2 phase separates less when UBA engages with Ub, in support of our prior observations whereby Ub abrogated UBQLN2 phase separation entirely (Dao et al., 2018). Our model also conforms with recent observations where UBQLN2 ΔUBA formed less puncta over time in neurons compared to FL and ΔUBL (Sharkey et al., 2018). Therefore, strong interactions involving the UBL or UBA domains outcompete the interdomain interactions within UBQLN2 to either promote or inhibit LLPS, respectively (Figure 5C). Importantly, our predictions address the effects of low-valency ligands on UBQLN2 LLPS. This is an important distinction as polyvalent ligands (e.g. long polyUb chains) may substantially alter the driving forces for UBQLN2 phase separation (Ruff et al., 2020).

## Conclusions

FL UBQLN2 appears poised to alter its phase behavior in either direction via interactions with binding partners. Interactions involving UBL- or UBA-binding partners, therefore, offer a rich and complex energetic landscape to modulate phase separation of UBQLN2 and other Ub-binding shuttle proteins (Yasuda et al., 2020). Additionally, our results suggest that client or chaperone proteins that engage with STI1 domains shift the balance of UBL/UBA interactions across the entire UBQLN2 protein (Hjerpe et al., 2016), thus impacting phase separation behavior. Although the presence of folded domains have been shown to alter the IDR-driven phase separation behavior of many proteins (FUS, hnRNPA1, LAF1 among others) (Burke et al., 2015; Molliex et al., 2015; Wei et al., 2017), our study and recent work by Martin and colleagues (Martin et al., 2021) highlight the need for detailed mapping of the interactions between folded domains and IDRs to further understand how cellular components affect phase-separating proteins and their functions. Our work also emphasizes the importance of identifying “stickers” and “spacers” within a protein system to understand how folded domains and intrinsically-disordered regions work together to modulate LLPS.

## Materials and Methods

### Subcloning, Protein Expression and Purification

The gene encoding human UBQLN2, a kind gift from Dr. Peter Howley’s laboratory (Addgene plasmid 8661), was subsequently cloned into a pET24b plasmid (Dao et al., 2018). Deletion constructs and mutations were created from this plasmid using the ThermoFisher Phusion Site-Directed Mutagenesis Kit and phosphorylated primers.

All UBQLN2 constructs, except isolated UBA and UBL, were transformed into Rosetta 2 (DE3) pLysS *E. coli* cells (Table S1). The cells were grown in LB media with final concentration of 50 mg/L kanamycin and 34 mg/L chloramphenicol to OD_600_ of 0.6-0.8, induced with 0.5 mM IPTG overnight, pelleted, frozen, then lysed using 20 mL of 20 mM NaPhosphate pH 6.8 buffer with 0.1 mg/mL DNAse, 5 mM MgCl_2,_ and 0.5 mM PMSF. The solution was mixed thoroughly using a pipette and centrifuged at 20,000 x g for 20 min. Supernatant was collected and warmed to room temperature. Solid NaCl was added to the solution to the final concentration of 0.5-1 M to induce phase separation, and the protein was then collected by centrifugation at 5000 x g for 20 min at room temperature. The protein pellet was then washed twice by re-solubilizing using cold, NaCl-free 20 mM NaPhosphate pH 6.8 buffer and re-precipitated by adding NaCl. In the final step, protein was solubilized again in 2 mL of cold, NaCl-free buffer, and the final product was obtained by running the solution through an 5 mL Hi-Trap (GE Healthcare) desalting column (Figure S1).

Isolated UBQLN2 UBA and UBL constructs were transformed into NiCo21 (DE3) *E. coli* cells. The cells were grown in LB with 50 mg/L kanamycin to OD_600_ of 0.6-0.8, induced with 0.5 mM IPTG overnight, pelleted, frozen, then resuspended in 20 mL of 20 mM NaPhosphate pH 7.2 buffer containing 0.1 mg/mL lysozyme, 0.1 mg/mL DNAse, 5 mM MgCl_2,_ and 0.5 mM PMSF. The resuspended cells were freeze-thawed for 3 rounds before centrifugation at 20,000 x g for 20 min. Supernatant was loaded onto a Ni-NTA column (GE Healthcare) equilibrated with the wash buffer (20 mM NaPhosphate pH 7.2 buffer containing 25 mM imidazole, and 0-100 mM NaCl). The column was washed with the wash buffer. The protein was then eluted with the elution buffer (20 mM NaPhosphate pH 7.2 buffer with 500 mM imidazole, and 0-100 mM NaCl), then dialyzed into 20 mM NaPhosphate pH 6.8 buffer containing 0.5 mM EDTA, 0.02% NaN_3_ overnight, concentrated, then further purified using a Superdex 75 HiLoad 16/600 column (GE Healthcare) equilibrated with 20 mM NaPhosphate pH 6.8, 0.5 mM EDTA and 0.02% NaN_3_.

### Temperature-dependent Spectrophotometric Turbidity Assays and Phase Diagrams

Protein samples were prepared using final concentrations of 5-80 μM protein in 20 mM NaPhosphate pH 6.8 buffer containing 0.5 mM EDTA, 0.02% NaN_3_, and 200 mM NaCl. Samples (400 μL) were then pipetted into quartz cuvettes (Starna Cells) on ice before loading into the Agilent Cary 3500 UV-Vis spectrophotometer. After incubating at 16°C for 2 minutes, OD_600_ values were recorded in 0.1°C increments as the samples were heated from 16°C to 60°C using a 1°C/minute temperature ramp rate. The OD_600_ versus temperature data were then fitted to a 4-parameter nonlinear regression curve y=*d*+(*a*-*d*)/[1+(x/*c*)^*b*] using MATLAB R2020a, where *a* and *d* are minimum and maximum absorbance values, *b* is the Hill slope reflecting steepness of the transition, and *c* is the temperature at the inflection point. Temperature-concentration phase diagrams were obtained by plotting the inflection point temperature (phase transition temperature) as a function of protein concentration. Plotted values are the average of at least four replicates (using at least two protein purifications) as described in figure legends.

Titration samples for turbidity assays were prepared using 10 μM UBQLN2 109-576 and 0-300 μM isolated UBL/UBA. Proteins were first mixed in 20 mM NaPhosphate pH 6.8 buffer containing 0.5 mM EDTA, 0.02% NaN_3_ on ice. Then a stock solution of 500 mM NaCl in 20 mM NaPhosphate pH 6.8, 0.5 mM EDTA and 0.02% NaN_3_ was added to a final NaCl concentration of 200 mM. Spectrophotometric measurements were carried out using the same method as described above.

### Fluorescence Microscopy

UBQLN2 constructs were prepared to contain 100 μM protein (spiked with Dylight-488 labeled protein, 1:100 ratio) in 20 mM NaPhosphate pH 6.8 buffer with 200 mM NaCl and 0.5 mM EDTA. Samples were added to concavity microscope slides (Eisco Labs) and covered with coverslips (MatTek) coated with 5% bovine serum albumin (BSA) to reduce rapid coating of protein droplets onto the coverslip, and incubated at 30 °C for 20 minutes. Phase separation was imaged on an ONI Nanoimager (Oxford Nanoimaging Ltd.) equipped with a Hamamatsu sCMOS ORCA flash 4.0 V3 camera using an Olympus 100 Å/1.4 N.A. objective. Images were prepared using Fiji (Schindelin et al., 2012) and FigureJ plugin (Mutterer and Zinck, 2013).

### NMR Spectroscopy

All NMR samples were prepared using ^15^N labeled protein (50-400 μM) in 20 mM NaPhosphate pH 6.8 buffer containing 0.5 mM EDTA, 0.02% NaN_3_, and 5% D_2_O. All spectra were collected using a Bruker Avance III 800 MHz NMR spectrometer equipped with a TCI cryoprobe at 25°C. The number of scans ranged from 8-64 depending on signal/noise ratio. All NMR data were processed using NMRPipe (Delaglio et al., 1995) and analyzed using CCPNMR Analysis 2.5 (Vranken et al., 2005) on NMRBox (Maciejewski et al., 2017). 2D ^1^H-^15^N TROSY-HSQC experiments were acquired using spectral widths of 12,019 and 2594.7 Hz in the direct ^1^H and indirect ^15^N dimensions, with 4806 and 240 total number of points, respectively, for ^1^H and ^15^N dimensions. Centers of frequency axes were 4.7 and 117 ppm for ^1^H and ^15^N, respectively. ^1^H-^15^N TROSY-HSQC spectra were processed and apodized using a Lorentz-to-Gauss window function in the ^1^H dimension, while ^15^N dimension was processed using a cosine squared bell function. Chemical shift perturbations (CSPs) were quantified as follows: 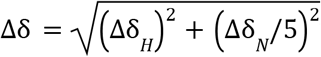 where Δδ_*H*_ and Δδ_*H*_ are the differences in ^1^H and ^15^N chemical shifts, respectively.

### NMR Titration Experiments

Unlabeled ligand was titrated into 200 μM samples of ^15^N UBL or UBA, and the binding was monitored by recording ^1^H-^15^N TROSY-HSQC or SOFAST-HMQC spectra as a function of ligand (L) concentration. At each titration point, we assumed that the chemical shift perturbation (CSP) for each backbone amide was a weighted average between the free (Δδ = 0) and ligand-bound (Δδ = Δδ_bound_) states, such that Δδ = Δδ_bound_ * [PL]/[P_total_]. Here, [PL] and [P_total_] represent the ligand-bound protein concentration, and P_total_ is the total protein concentration of the ^15^N-labelled protein. Data fitting for each amide was performed using an in-house MATLAB program, assuming a single-site (1:1 stoichiometry) binding model. Only residues with CSP > 0.03 ppm at the titration endpoint were considered for *K*_d_ determination. Reported *K*_d_ values were averages of residue-specific *K*_d_ values with errors reflecting standard deviation of these values.

### Resonance assignments for UBQLN2 UBL (1-107)

To obtain chemical shift assignments, NMR samples consisting of 500 mM ^15^N,^13^C-labeled UBL were prepared in pH 6.8 buffer (see above) with 20 mM NaPhosphate, 0.5 mM EDTA, 0.02% w/v NaN_3_, and 5% D_2_O. All experiments were collected at 298 K. Chemical shift assignments of the backbone resonances (H^N^, N, Ca, and CΔ) were obtained using 2D ^1^H-^15^N, HSQC, and triple resonance experiments (HNCACB, CBCACONH, and HNN). Acquisition times for the triple resonance experiments were 80 ms in the direct ^1^H dimensions, 18 ms in the indirect ^15^N dimensions, and 6 ms in the indirect Ca/CΔ dimensions. Spectral widths were 16 ppm in ^1^H^N^, 68 ppm in ^13^Ca/Cb dimension, 27 ppm in indirect ^15^N. Non-uniform sampling (NUS) was employed for all triple resonance experiments. Experiments were acquired with 25% (CBCACONH, HNCACB) or 30% (HNN) sampling using the Poisson Gap sampling method (Hyberts et al., 2010). NUS spectra were processed using SMILE and NMRPipe (Delaglio et al., 1995; Ying et al., 2017) and employed standard apodization parameters and linear prediction in the indirect dimensions. Using these experiments, we successfully assigned amide backbone resonances (H^N^, N) for 99% (97/98 assignable) of all residues.

## Abbreviations and symbols

c_sat_: Saturation concentration
CSP: Chemical shift perturbation
DSK2: Ubiquitin domain-containing protein DSK2
FUS: Fused-in-sarcoma
FL: Full-length
hnRNPA1: Heterogeneous nuclear ribonucleoprotein A1
HSP70: Heat shock protein 70
HSQC: Heteronuclear single quantum coherence
IDR: Intrinsically disordered region
*K*_d_: Binding dissociation constant
LLPS: Liquid-liquid phase separation
NMR: Nuclear magnetic resonance
PRE: Paramagnetic relaxation enhancement
PQC: Protein quality control
PXX: Proline-rich region
STI1: Stress induced protein 1 (STI1)-like domain
TROSY: Transverse-relaxation optimized spectroscopy
UBA: Ubiquitin-associating domain
UBL: Ubiquitin-like domain
UBQLN2: Ubiquilin-2
Ub: Ubiquitin
UPS: Ubiquitin-proteasome system

## Acknowledgements

Initial UBL and UBA turbidity experiments with 109-576 were supported by NSF CAREER 1750462 to C.A.C.. The construction of the 109-576 library, NMR experiments involving the 109-576 library, design of UBL and UBA mutants, and subsequent analyses were supported by NIH R01 GM136946 to C.A.C.. Data collection with a Bruker Ascend 800 MHz NMR magnet was supported by NIH shared instrumentation grant 1S10OD012254. T.Z. was partially supported by the Kathy Walters fund. We acknowledge Dr. Thuy Dao for assistance in protein purification, molecular cloning and collecting fluorescence microscopy images, and Yiran Yang for assistance in protein purification and NMR spectra collection. We thank the past and present members of the Castañeda lab, especially Dr. Thuy Dao and Dr. Sarasi Galagedera, for insightful discussions and comments throughout this project.

## Supplementary Material

Supplementary material includes eight figures, five tables, and four movies.

## References

Alexander, E.J., Ghanbari Niaki, A., Zhang, T., Sarkar, J., Liu, Y., Nirujogi, R.S., Pandey, A., Myong, S., and Wang, J. (2018). Ubiquilin 2 modulates ALS/FTD-linked FUS-RNA complex dynamics and stress granule formation. Proc. Natl. Acad. Sci. 115, E11485–E11494.

Bjellqvist, B., Hughes, G.J., Pasquali, C., Paquet, N., Ravier, F., Sanchez, J.C., Frutiger, S., and Hochstrasser, D. (1993). The focusing positions of polypeptides in immobilized pH gradients can be predicted from their amino acid sequences. Electrophoresis 14, 1023–1031.

Burke, K.A., Janke, A.M., Rhine, C.L., and Fawzi, N.L. (2015). Residue-by-Residue View of In Vitro FUS Granules that Bind the C-Terminal Domain of RNA Polymerase II. Mol. Cell 60, 231–241.

Chen, L., Shinde, U., Ortolan, T.G., and Madura, K. (2001). Ubiquitin-associated (UBA) domains in Rad23 bind ubiquitin and promote inhibition of multi-ubiquitin chain assembly. EMBO Rep. 2, 933–938.

Chen, X., Randles, L., Shi, K., Tarasov, S.G., Aihara, H., and Walters, K.J. (2016). Structures of Rpn1 T1:Rad23 and hRpn13:hPLIC2 Reveal Distinct Binding Mechanisms between Substrate Receptors and Shuttle Factors of the Proteasome. Structure 24, 1257–1270.

Chen, X., Ebelle, D.L., Wright, B.J., Sridharan, V., Hooper, E., and Walters, K.J. (2019). Structure of hRpn10 Bound to UBQLN2 UBL Illustrates Basis for Complementarity between Shuttle Factors and Substrates at the Proteasome. J. Mol. Biol. 431, 939–955.

Dao, T.P., and Castañeda, C.A. (2020). Ubiquitin-Modulated Phase Separation of Shuttle Proteins: Does Condensate Formation Promote Protein Degradation? BioEssays 42, 2000036.

Dao, T.P., Kolaitis, R.-M., Kim, H.J., O’Donovan, K., Martyniak, B., Colicino, E., Hehnly, H., Taylor, J.P., and Castañeda, C.A. (2018). Ubiquitin Modulates Liquid-Liquid Phase Separation of UBQLN2 via Disruption of Multivalent Interactions. Mol. Cell 69, 965–978.e6.

Dao, T.P., Martyniak, B., Canning, A.J., Lei, Y., Colicino, E.G., Cosgrove, M.S., Hehnly, H., and Castañeda, C.A. (2019). ALS-Linked Mutations Affect UBQLN2 Oligomerization and Phase Separation in a Position- and Amino Acid-Dependent Manner. Structure 27, 937–951.e5.

Delaglio, F., Grzesiek, S., Vuister, G.W., Zhu, G., Pfeifer, J., and Bax, A. (1995). NMRPipe: a multidimensional spectral processing system based on UNIX pipes. J. Biomol. NMR 6, 277–293.

Díaz-Martínez, L.A., Kang, Y., Walters, K.J., and Clarke, D.J. (2006). Yeast UBL-UBA proteins have partially redundant functions in cell cycle control. Cell Div. 1, 28.

Heir, R., Ablasou, C., Dumontier, E., Elliott, M., Fagotto-Kaufmann, C., and Bedford, F.K. (2006). The UBL domain of PLIC-1 regulates aggresome formation. EMBO Rep. 7, 1252–1258.

Hjerpe, R., Bett, J.S., Keuss, M.J., Solovyova, A., McWilliams, T.G., Johnson, C., Sahu, I., Varghese, J., Wood, N., Wightman, M., et al. (2016). UBQLN2 Mediates Autophagy-Independent Protein Aggregate Clearance by the Proteasome. Cell 166, 935–949.

Hyberts, S.G., Takeuchi, K., and Wagner, G. (2010). Poisson-Gap Sampling and Forward Maximum Entropy Reconstruction for Enhancing the Resolution and Sensitivity of Protein NMR Data. J. Am. Chem. Soc. 132, 2145–2147.

Jones, D.T., and Cozzetto, D. (2015). DISOPRED3: precise disordered region predictions with annotated protein-binding activity. Bioinformatics 31, 857–863.

Kjaergaard, M., Glavina, J., and Chemes, L.B. (2021). Chapter Six - Predicting the effect of disordered linkers on effective concentrations and avidity with the “Ceff calculator” app. In Methods in Enzymology, M. Merkx, ed. (Academic Press), pp. 145–171.

Ko, H.S., Uehara, T., Tsuruma, K., and Nomura, Y. (2004). Ubiquilin interacts with ubiquitylated proteins and proteasome through its ubiquitin-associated and ubiquitin-like domains. FEBS Lett. 566, 110–114.

Kurlawala, Z., Shah, P.P., Shah, C., and Beverly, L.J. (2017). The STI and UBA domains of UBQLN1 are critical determinants of substrate interaction and proteostasis. J. Cell. Biochem. 118, 2261–2270.

Lee, D.Y., Arnott, D., and Brown, E.J. (2013). Ubiquilin4 is an adaptor protein that recruits Ubiquilin1 to the autophagy machinery. EMBO Rep. 14, 373–381.

Lowe, E.D., Hasan, N., Trempe, J.-F., Fonso, L., Noble, M.E.M., Endicott, J.A., Johnson, L.N., and Brown, N.R. (2006). Structures of the Dsk2 UBL and UBA domains and their complex. Acta Crystallogr. D Biol. Crystallogr. 62, 177–188.

Maciejewski, M.W., Schuyler, A.D., Gryk, M.R., Moraru, I.I., Romero, P.R., Ulrich, E.L., Eghbalnia, H.R., Livny, M., Delaglio, F., and Hoch, J.C. (2017). NMRbox: A Resource for Biomolecular NMR Computation. Biophys. J. 112, 1529–1534.

Martin, E.W., Thomasen, F.E., Milkovic, N.M., Cuneo, M.J., Grace, C.R., Nourse, A., Lindorff-Larsen, K., and Mittag, T. (2021). Interplay of folded domains and the disordered low-complexity domain in mediating hnRNPA1 phase separation. BioRxiv 2020.05.15.096966.

Martinez-Fonts, K., Davis, C., Tomita, T., Elsasser, S., Nager, A.R., Shi, Y., Finley, D., and Matouschek, A. (2020). The proteasome 19S cap and its ubiquitin receptors provide a versatile recognition platform for substrates. Nat. Commun. 11, 477.

Molliex, A., Temirov, J., Lee, J., Coughlin, M., Kanagaraj, A.P., Kim, H.J., Mittag, T., and Taylor, J.P. (2015). Phase separation by low complexity domains promotes stress granule assembly and drives pathological fibrillization. Cell 163, 123–133.

Mutterer, J., and Zinck, E. (2013). Quick-and-clean article figures with FigureJ. J. Microsc. 252, 89–91.

Nguyen, K., Puthenveetil, R., and Vinogradova, O. (2017). Investigation of the adaptor protein PLIC-2 in multiple pathways. Biochem. Biophys. Rep. 24055808 9.

Ohno, A., Jee, J., Fujiwara, K., Tenno, T., Goda, N., Tochio, H., Kobayashi, H., Hiroaki, H., and Shirakawa, M. (2005). Structure of the UBA Domain of Dsk2p in Complex with Ubiquitin: Molecular Determinants for Ubiquitin Recognition. Structure 13, 521–532.

Ruff, K.M., Roberts, S., Chilkoti, A., and Pappu, R.V. (2018). Advances in Understanding Stimulus-Responsive Phase Behavior of Intrinsically Disordered Protein Polymers. J. Mol. Biol. 430, 4619–4635.

Ruff, K.M., Dar, F., and Pappu, R.V. (2020). Ligand Effects on Phase Separation of Multivalent Macromolecules. BioRxiv 2020.08.15.252346.

Schindelin, J., Arganda-Carreras, I., Frise, E., Kaynig, V., Longair, M., Pietzsch, T., Preibisch, S., Rueden, C., Saalfeld, S., Schmid, B., et al. (2012). Fiji: an open-source platform for biological-image analysis. Nat. Methods 9, 676–682.

Sharkey, L.M., Safren, N., Pithadia, A.S., Gerson, J.E., Dulchavsky, M., Fischer, S., Patel, R., Lantis, G., Ashraf, N., Kim, J.H., et al. (2018). Mutant UBQLN2 promotes toxicity by modulating intrinsic self-assembly. Proc. Natl. Acad. Sci. 115, E10495–E10504.

Sun, D., Wu, R., Zheng, J., Li, P., and Yu, L. (2018). Polyubiquitin chain-induced p62 phase separation drives autophagic cargo segregation. Cell Res. 28, 405–415.

Tse, M.K., Hui, S.K., Yang, Y., Yin, S.-T., Hu, H.-Y., Zou, B., Wong, B.C.Y., and Sze, K.H. (2011). Structural Analysis of the UBA Domain of X-linked Inhibitor of Apoptosis Protein Reveals Different Surfaces for Ubiquitin-Binding and Self-Association. PLOS ONE 6, e28511.

Urry, D.W., Gowda, D.C., Parker, T.M., Luan, C.H., Reid, M.C., Harris, C.M., Pattanaik, A., and Harris, R.D. (1992). Hydrophobicity scale for proteins based on inverse temperature transitions. Biopolymers 32, 1243–1250.

Vranken, W.F., Boucher, W., Stevens, T.J., Fogh, R.H., Pajon, A., Llinas, M., Ulrich, E.L., Markley, J.L., Ionides, J., and Laue, E.D. (2005). The CCPN data model for NMR spectroscopy: Development of a software pipeline. Proteins Struct. Funct. Bioinforma. 59, 687–696.

Walters, K.J., Kleijnen, M.F., Goh, A.M., Wagner, G., and Howley, P.M. (2002). Structural studies of the interaction between ubiquitin family proteins and proteasome subunit S5a. Biochemistry 41, 1767–1777.

Wei, M.-T., Elbaum-Garfinkle, S., Holehouse, A.S., Chen, C.C.-H., Feric, M., Arnold, C.B., Priestley, R.D., Pappu, R.V., and Brangwynne, C.P. (2017). Phase behaviour of disordered proteins underlying low density and high permeability of liquid organelles. Nat. Chem. 9, 1118–1125.

Xue, B., Dunbrack, R.L., Williams, R.W., Dunker, A.K., and Uversky, V.N. (2010). PONDR-FIT: A Meta-Predictor of Intrinsically Disordered Amino Acids. Biochim. Biophys. Acta 1804, 996–1010.

Yang, Y., Jones, H.B., Dao, T.P., and Castañeda, C.A. (2019). Single Amino Acid Substitutions in Stickers, but Not Spacers, Substantially Alter UBQLN2 Phase Transitions and Dense Phase Material Properties. J. Phys. Chem. B 123, 3618–3629.

Yasuda, S., Tsuchiya, H., Kaiho, A., Guo, Q., Ikeuchi, K., Endo, A., Arai, N., Ohtake, F., Murata, S., Inada, T., et al. (2020). Stress- and ubiquitylation-dependent phase separation of the proteasome. Nature 578, 296–300.

Ying, J., Delaglio, F., Torchia, D.A., and Bax, A. (2017). Sparse multidimensional iterative lineshape-enhanced (SMILE) reconstruction of both non-uniformly sampled and conventional NMR data. J. Biomol. NMR 68, 101–118.

Zaffagnini, G., Savova, A., Danieli, A., Romanov, J., Tremel, S., Ebner, M., Peterbauer, T., Sztacho, M., Trapannone, R., Tarafder, A.K., et al. (2018). p62 filaments capture and present ubiquitinated cargos for autophagy. EMBO J. 37, e98308.

Zhang, D., Raasi, S., and Fushman, D. (2008). Affinity Makes the Difference: Nonselective Interaction of the UBA Domain of Ubiquilin-1 with Monomeric Ubiquitin and Polyubiquitin Chains. J. Mol. Biol. 377, 162–180.

Zheng, T., Yang, Y., and Castañeda, C.A. (2020). Structure, dynamics and functions of UBQLNs: at the crossroads of protein quality control machinery. Biochem. J. 477, 3471–3497.

Zhou, H.-X., Nguemaha, V., Mazarakos, K., and Qin, S. (2018). Why Do Disordered and Structured Proteins Behave Differently in Phase Separation? Trends Biochem. Sci. 43, 499–516.

Zientara-Rytter, K., and Subramani, S. (2019). The Roles of Ubiquitin-Binding Protein Shuttles in the Degradative Fate of Ubiquitinated Proteins in the Ubiquitin-Proteasome System and Autophagy. Cells 8.

