## Supplementary Material for "Previously uncharacterized interactions between the folded and intrinsically disordered domains impart asymmetric effects on UBQLN2 phase separation"

Supplementary Figures S1-S8
Supplementary Tables S1-S5
Supplementary Movies S1-S4

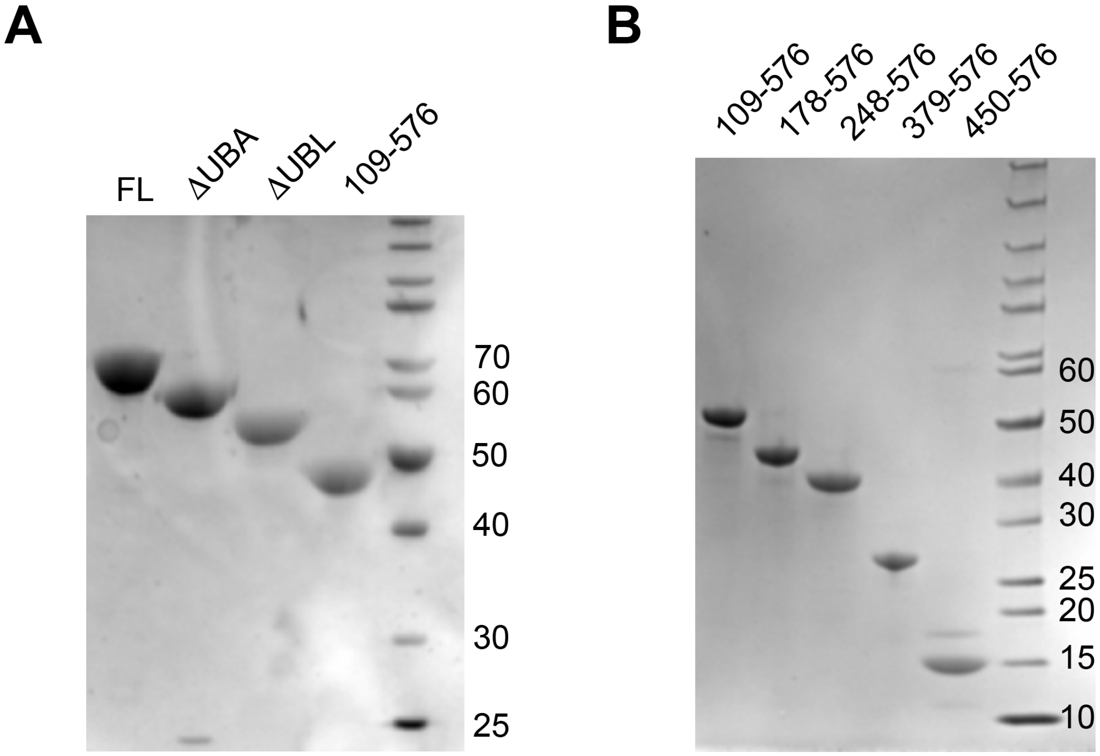


**Supplementary Figure S1.** SDS-PAGE gels of UBQLN2 constructs used in this study.

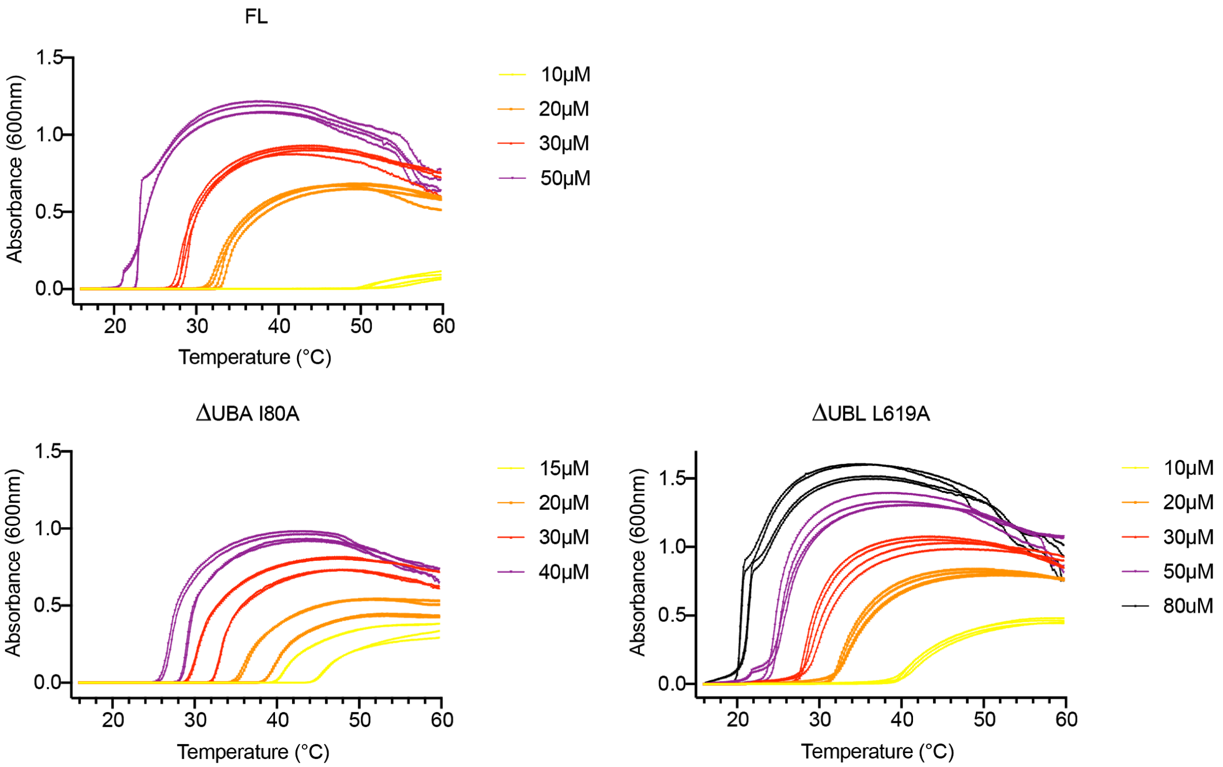


**Supplementary Figure S2.** Representative individual spectrophotometric turbidity experiments used to construct temperature-concentration phase diagrams in Figures 1 and 4. For each protein concentration, turbidity experiments were repeated using protein from at least two protein purifications for a total of at least four trials.

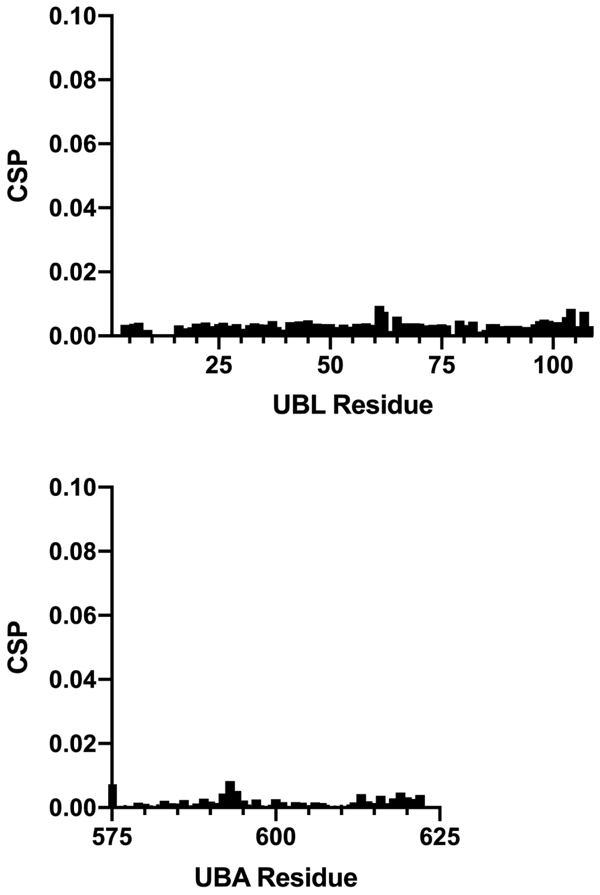


**Supplementary Figure S3.** Little or no concentration dependence of the isolated UBL and UBA domains. CSPs of UBL or UBA backbone amide resonances were measured between NMR spectra collected using protein concentrations of 50 to 400 μM.

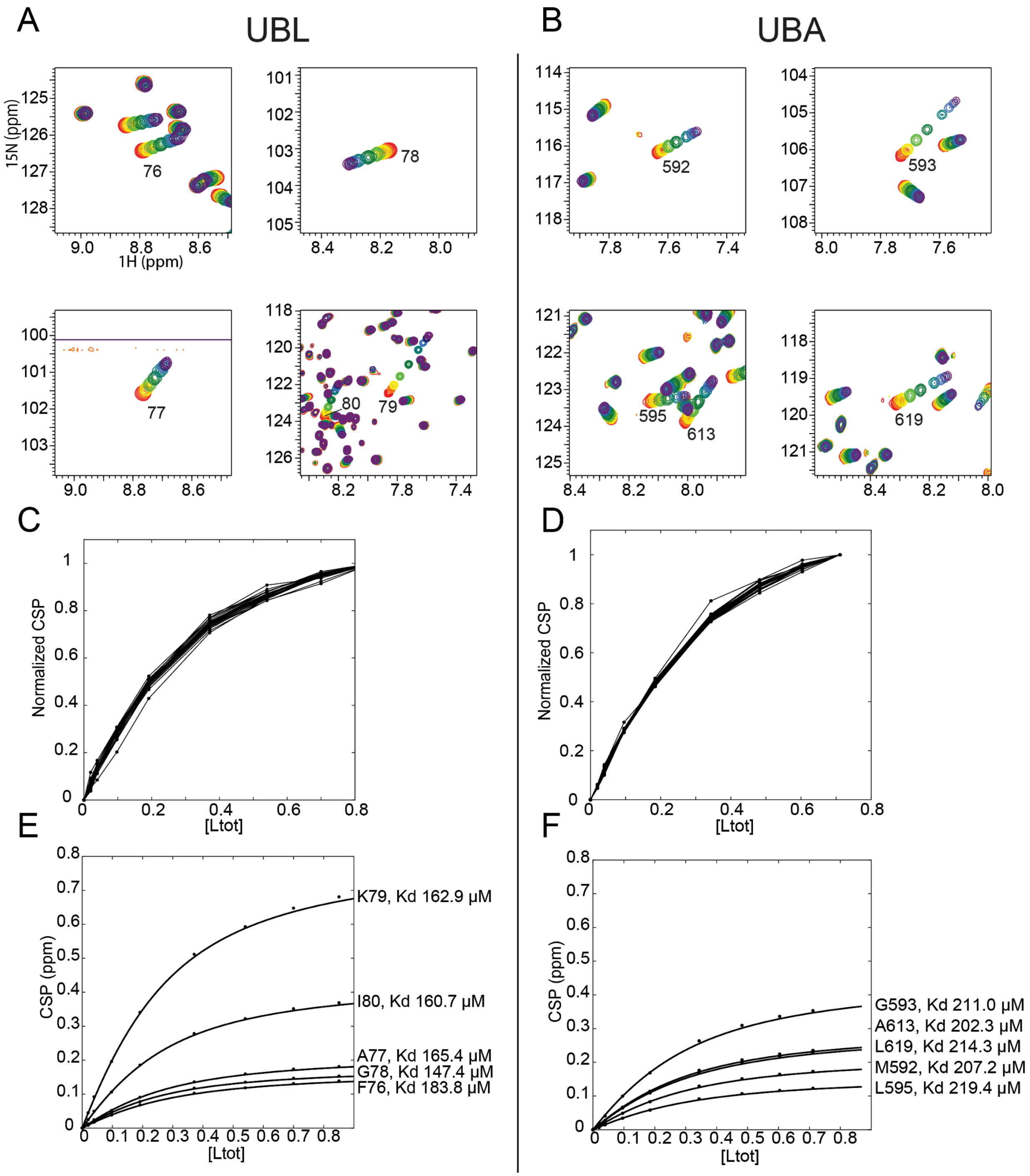


**Supplementary Figure S4.** NMR titration data from $^{15}N$-$^{1}H$ HSQC spectra for UBQLN2 UBL:UBA interactions. Representative amide resonance trajectories for (A) UBL or (B) UBA residues as UBA or UBL were added, respectively. (C, D) Normalized CSPs were plotted against total ligand concentrations for (C) UBL and (D) UBA resonances. (E, F) A single-site binding model (see Methods) was fit to the titration CSP data (black circles) to determine residue-specific $K_d$. Curve fits for representative residues in the (E) UBL and (F) UBA domains are shown here.

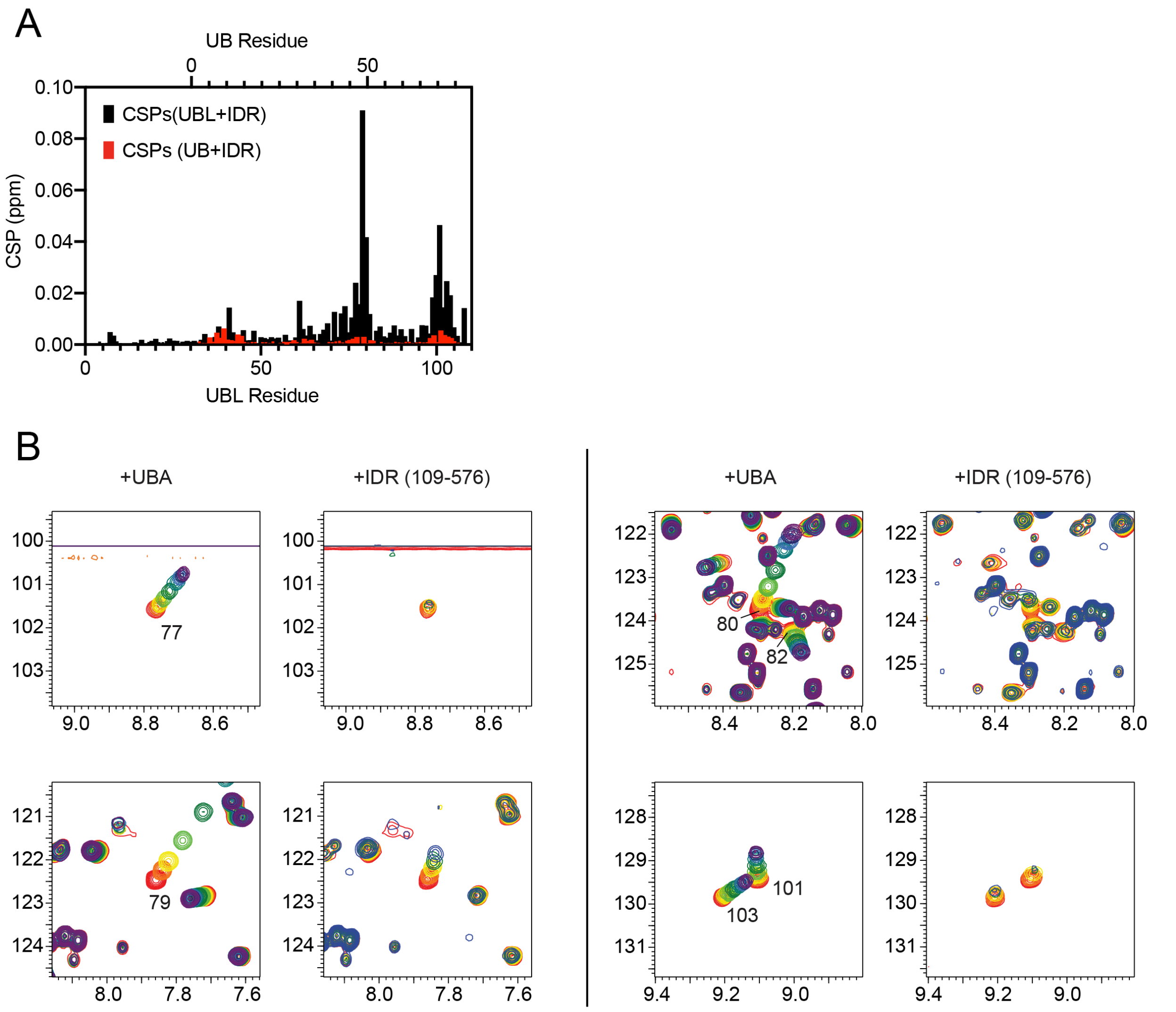


**Supplementary Figure S5.** Analysis of UBL interactions with 109-576 ($\Delta$UBL$\Delta$UBA). (A) Comparison of amide trajectories from $^{1}$H-$^{15}$N HSQC spectra for UBL residues 77, 79, 80, 82, 101, and 103 as a function of UBA or 109-576 titrations. The UBA titration was performed using 200 μM $^{15}$N UBL, whereas the 109-576 titration was collected using 50 μM $^{15}$N UBL. UBL resonances broadened extensively as 109-576 was titrated, unlike in the presence of UBA. UBA concentrations were 0 μM (red), 20 μM (orange), 40 μM (yellow), 100μM (lime), 190 μM (green), 370 μM (teal), 540 μM (sky blue), 700 μM (blue), 850 μM (purple). 109-576 concentrations were 0 μM (red), 10 μM (orange), 30 μM (yellow), 50 μM (green), 70 μM (blue). (B) Comparison of UBL CSPs in equimolar UBL:109-576 mixture versus Ub CSPs in equimolar Ub:109-576 mixture (each protein at 50 μM). UBL and Ub sequences were structurally aligned and corresponding sequence alignment used for x-axes (top: Ub, bottom: UBL).

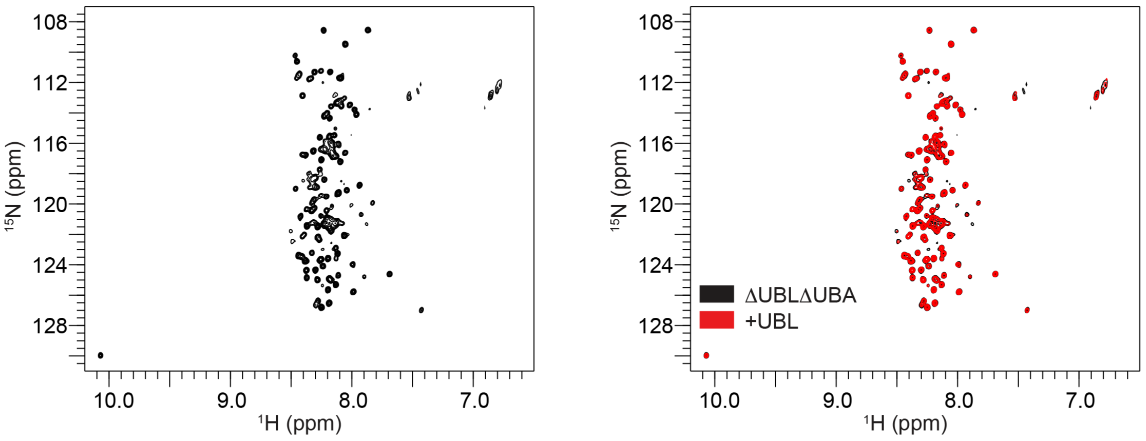


**Supplementary Figure S6.** $^{1}$H-$^{15}$N TROSY-HSQC NMR spectra of 50 µM UBQLN2 109-576 (ΔUBLΔUBA) in the absence (left) and presence (right) of equimolar amounts of UBL (1-107).

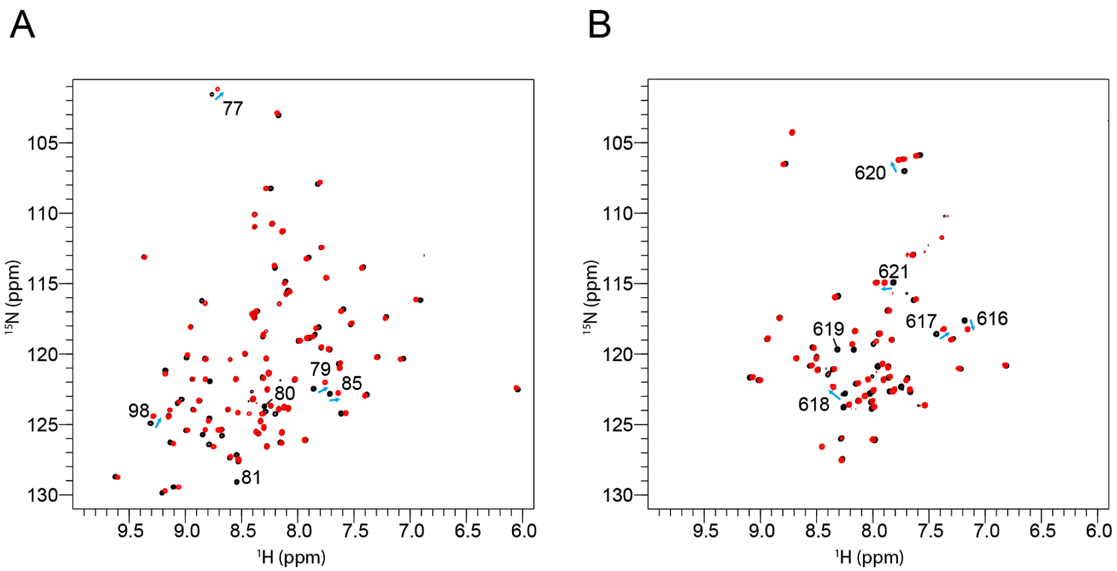


**Supplementary Figure S7.** Comparison of $^{1}$H-$^{15}$N TROSY-HSQC NMR spectra of (A) 50 µM WT UBL (black) and I80A UBL (red), (B) 200 µM WT UBA (black) and L619A UBA (red). Amide resonances that exhibited significant CSPs were highlighted (blue arrows), as well as residues I80 and L619.

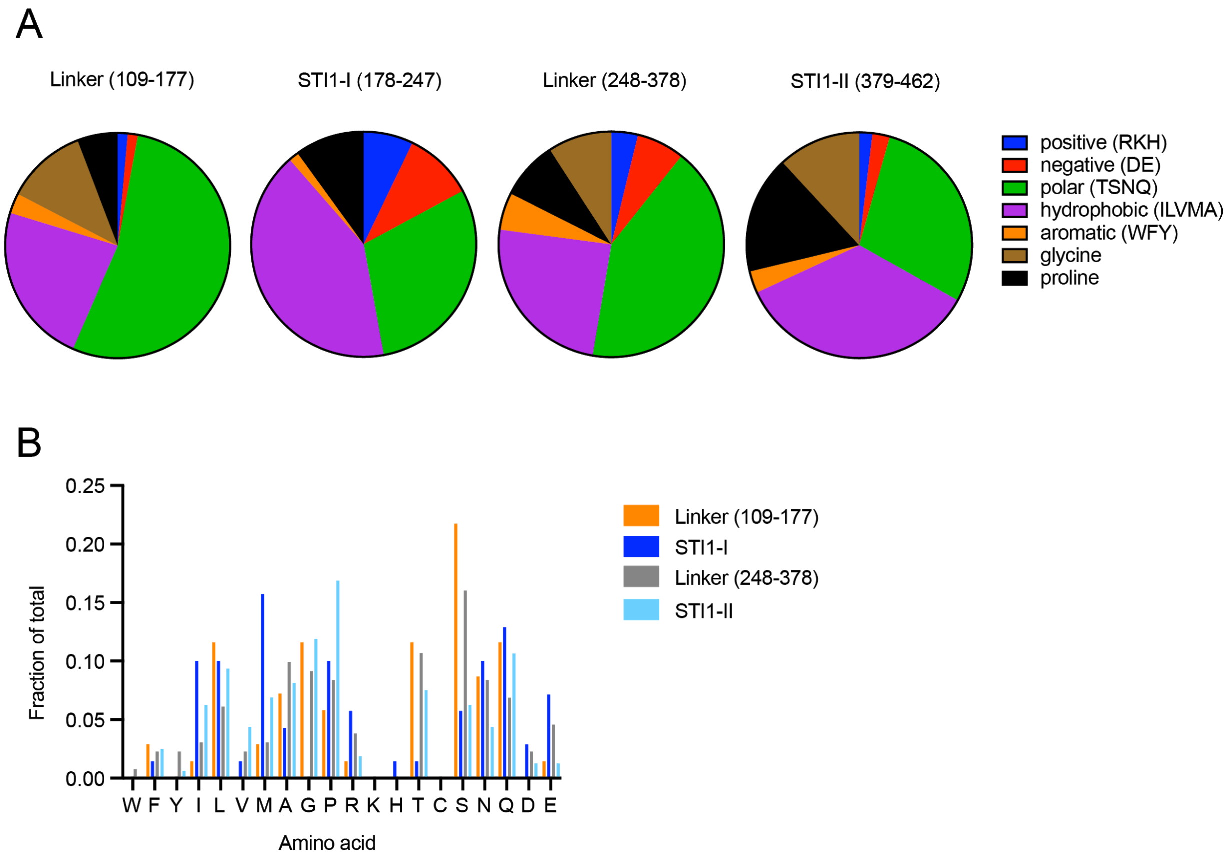


**Supplementary Figure S8.** Amino acid composition for linkers and domains within residues 109-576 of UBQLN2. (A) Pie chart follows the general organization in (Ruff et al., 2018). (B) Fraction of amino acids in each color-coded region within 109-576. Amino acids ranked in decreasing hydrophobicity from left to right, according to scale determined by (Urry et al., 1992).

**Supplementary Table S1.** UBQLN2 Constructs used for this study.

| UBQLN2 Construct | Residues | Extinction Coefficient ($M^{-1}$ $cm^{-1}$) | Special Notes |
|---|---|---|---|
| FL | 1-624 | 11460 | Tagless |
| ΔUBA | 1-576 | 11460 | Tagless |
| ΔUBL | 109-624 | 11460 | Tagless |
| 109-576 (ΔUBLΔUBA) | 109-576 | 11460 | Tagless |
| 178-576 | 178-576 | 11460 | Tagless |
| 248-576 | 248-576 | 11460 | Tagless |
| 379-576 | 379-576 | 1490 | Tagless |
| 450-576 | 450-576 | 0 | Tagless, concentration measured as described in Methods |
| UBL | 1-107 | 5500 | C-terminal WHHHHHH |
| UBA | 571-624 | 5500 | C-terminal WHHHHHH |
| I80A UBL | 1-107 | 5500 | C-terminal WHHHHHH |
| L619A UBA | 571-624 | 5500 | C-terminal WHHHHHH |
| I80A ΔUBA | 1-576 | 11460 | Tagless |
| L619A ΔUBL | 109-624 | 11460 | Tagless |

**Supplementary Table S2.** UBL:UBA binding affinity measurements.

| | $K_d$ |
|---|---|
| $^{15}$N UBL + unlabeled UBA | 178 ± 41 μM |
| $^{15}$N UBA + unlabeled UBL | 210.1 ± 15.3 μM |
| $^{15}$N UBA + unlabeled UBL (Nguyen et al., 2017) | 175 ± 25 μM |

**Supplementary Table S3.** Residue-by-residue $K_d$ determination from NMR titration curves.

| Residue number | Max CSP (ppm) | $K_d$ (mM) |
|---|---|---|
| **UBL** | | |
| 79 | 0.6809 | 0.1629 |
| 80 | 0.3691 | 0.1607 |
| 77 | 0.1810 | 0.1654 |
| 78 | 0.1532 | 0.1474 |
| 76 | 0.1387 | 0.1838 |
| 81 | 0.1378 | 0.1713 |
| 104 | 0.1377 | 0.1645 |
| 101 | 0.1290 | 0.1583 |
| 99 | 0.1056 | 0.1789 |
| 82 | 0.1016 | 0.1677 |
| **UBA** | | |
| 593 | 0.3530 | 0.2110 |
| 613 | 0.2352 | 0.2023 |
| 619 | 0.2291 | 0.2143 |
| 592 | 0.1729 | 0.2072 |
| 595 | 0.1229 | 0.2194 |
| 594 | 0.1030 | 0.1663 |
| 616 | 0.0948 | 0.2073 |
| 590 | 0.0781 | 0.2170 |
| 620 | 0.0748 | 0.2336 |
| 615 | 0.0695 | 0.2096 |

**Supplementary Table S4.** UBQLN2 equivalent of highlighted DSK2 residues in Figure 2C.

| UBL residues | | UBA residues | |
|---|---|---|---|
| UBQLN2 equivalent | DSK2 | UBQLN2 equivalent | DSK2 |
| F76 | Y46 | M592 | M342 |
| A77 | S47 | G593 | G343 |
| G78 | G48 | L595 | F345 |
| K79 | K49 | A613 | G363 |
| I80 | I50 | E616 | D366 |
| L81 | L51 | L619 | L369 |
| V101 | V71 | | |

**Supplementary Table S5.** Calculated isoelectric point (pI) of UBQLN2 constructs.

| Construct name | Calculated pI (Bjellqvist et al., 1993) |
|---|---|
| FL | 5.15 |
| UBL (1-107) | 9.35 |
| UBA (577-624) | 4.59 |
| 109-576 | 4.54 |
| 178-576 | 4.50 |
| 248-576 | 4.39 |
| 379-576 | 4.36 |
| 450-576 | 4.25 |

**Supplementary Movies**

**Movies S1-S4:** UBQLN2 droplets are liquid-like. Solutions of UBQLN2 domain deletion constructs under identical experimental conditions as shown in Figure 1. Short movies (between 10 and 30 seconds) illustrate UBQLN2 droplets fusing with each other, indicative of their liquid-like properties. Scale bar is 2 or 5 μm, as indicated in the movies.
